# Quantifying brain connectivity signatures by means of polyconnectomic scoring

**DOI:** 10.1101/2023.09.26.559327

**Authors:** Ilan Libedinsky, Koen Helwegen, Laura Guerrero Simón, Marius Gruber, Jonathan Repple, Tilo Kircher, Udo Dannlowski, Martijn P. van den Heuvel

## Abstract

A broad range of neuropsychiatric disorders are associated with alterations in macroscale brain circuitry and connectivity. Identifying consistent brain patterns underlying these disorders by means of structural and functional MRI has proven challenging, partly due to the vast number of tests required to examine the entire brain, which can lead to an increase in missed findings. In this study, we propose polyconnectomic score (PCS) as a metric designed to quantify the presence of disease-related brain connectivity signatures in connectomes. PCS summarizes evidence of brain patterns related to a phenotype across the entire landscape of brain connectivity into a subject-level score. We evaluated PCS across four brain disorders (autism spectrum disorder, schizophrenia, attention deficit hyperactivity disorder, and Alzheimer’s disease) and 14 studies encompassing ∼35,000 individuals. Our findings consistently show that patients exhibit significantly higher PCS compared to controls, with effect sizes that go beyond other single MRI metrics ([min, max]: Cohen’s *d* = [0.30, 0.87], *AUC* = [0.58, 0.73]). We further demonstrate that PCS serves as a valuable tool for stratifying individuals, for example within the psychosis continuum, distinguishing patients with schizophrenia from their first-degree relatives (*d* = 0.42, *p* = 4 x 10^−3^, FDR-corrected), and first-degree relatives from healthy controls (*d* = 0.34, *p* = 0.034, FDR-corrected). We also show that PCS is useful to uncover associations between brain connectivity patterns related to neuropsychiatric disorders and mental health, psychosocial factors, and body measurements.

## Introduction

Understanding the circuitry architecture of the human brain is a major goal in neuroscience and medicine. Magnetic resonance imaging (MRI) provides a non-invasive method for mapping macroscale structural and functional connections in the brain. The field of network neuroscience^1^ describes and analyzes the intricate system of connections within the nervous system, examining variations in brain connectivity underlying healthy and pathological conditions^2,3^.

Detecting consistent and reliable neurobiological processes underlying neuropsychiatric disorders^4,5^ has proven challenging due to methodological limitations^6–9^ and the inherent heterogeneity of the phenotypes studied^4,10^, among other factors^11,12^. One methodological challenge arises when examining the extensive number of connections in the brain. Network studies often overlook the contribution from connections that do not reach significance after controlling for multiple comparisons^9,13^, and these ‘missed connections’^6^ might constitute a substantial portion of the truly involved brain circuitry^14–16^. The search for brain signatures underlying neuropsychiatric disorders is further complicated by the co-occurrence of different disorders within a single individual^17^ and the overlapping neurobiological alterations among conditions^18–22^. Studying heterogeneous populations can result in a spatially distributed neural circuitry associated with a phenotype, complicating the identification of consistent brain patterns across individuals.

We propose polyconnectomic score (PCS)^3^ as a metric to capture connectome signatures in individual subjects. Drawing inspiration from polygenic score in genetics^23^, PCS quantifies the presence of brain circuitry associated with a specific phenotype in an individual’s brain connectome. By distilling the biological evidence of both subtle and pronounced connectivity alterations into a single score, PCS provides a global depiction of the disorder-related neural circuitry present in an individual while reducing the number of tests required. We show the utility of PCS in three different applications: identifying individuals predisposed to disease based on their brain connectivity, stratifying patients according to disease liability, and uncovering brain-behavior correlations. PCS provides a way to quantify the presence of connectivity signatures related to a phenotype within a connectome, facilitating further investigation into the brain’s role in health and disease states.

## Materials and Methods

### Studies and subjects

Resting-state functional MRI (fMRI) data from a total of 34,570 individuals were included from 14 different studies. Each study received approval from the relevant ethics committee, and participants provided written informed consent. Descriptions of these studies and their corresponding scanning parameters are detailed in the Supplemental Methods and Table S1, respectively.

We initially evaluated case-control differences in PCS by examining 12 studies including 4,610 healthy controls and 2,683 individuals with schizophrenia (SCZ), autism spectrum disorder (ASD), attention deficit hyperactivity disorder (ADHD), or Alzheimer’s disease (AD)^24–35^. Table 1 provides a demographic overview of these studies. Additionally, a 13th study was included for stratification of individuals based on their liability to psychosis, comprising individuals with SCZ, schizoaffective disorder (SCA), and psychotic bipolar disorder (BD; *n* = 126, 59, and 72, respectively), as well as their first-degree relatives (*n* = 113, 71, and 75, respectively), and healthy controls (*n* = 88)^36^. We further incorporated data from the UK Biobank (UKB)^37^ to measure brain-behavior associations, including 26,673 individuals with both neuroimaging and behavioral assessments.

**Table 1.**
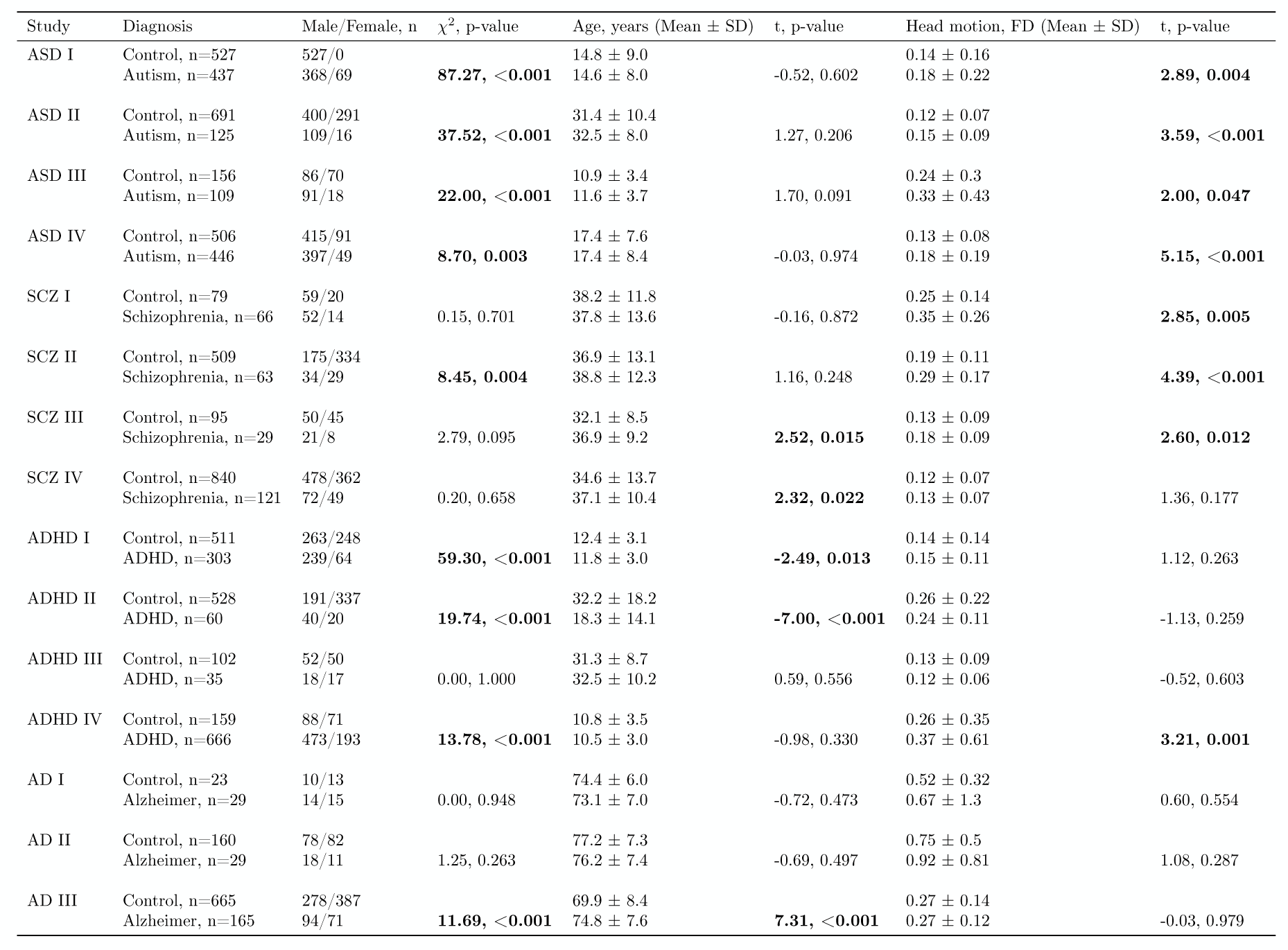
Demographic overview of included studies. A demographic overview of the studies included in the main analysis is presented. For each study (first column), the table provides the number of patients and controls (second column), the number of males and females (third column), the mean and standard deviation for age (fifth column) and head motion (seventh column). Statistics used to evaluate differences in these measurements between patients and controls are also given (fourth, sixth, and eighth columns). A positive *t*-value indicates higher values in patients compared to controls. Bolded values denote statistically significant group differences. *ASD*, autism spectrum disorder; *SCZ*, schizophrenia; *ADHD*, attention deficit hyperactivity disorder; *AD*, Alzheimer’s disease; *FD*, framewise displacement; *SD*, standard deviation.

### Data processing

Anatomical T1-weighted images were parcellated into 68 cortical regions following the Desikan–Killiany atlas^38^ using FreeSurfer v7.1.1^39^. Functional connectivity (FC) reconstruction was done with CATO 3.1.2^20^. Motion parameters and the signal intensity of white matter and cerebrospinal fluid were regressed out from the fMRI time-series. Global mean correction was performed by regressing out the mean signal intensity of all voxels in the brain from the time-series^40^. Next, bandpass filtering (0.01 - 0.1 Hz) and motion scrubbing^41^ (max FD = 0.25, max DVARS = 1.5) were applied to the BOLD time-series. FC between all pairs of brain regions was estimated by extracting the mean time-series from the cortical regions and computing Pearson’s correlation coefficient between each pair of regions^42,43^. Quality control was performed based on two criteria: individuals were removed if either their mean positive connections or more than 1% of their total connections deviated beyond three standard deviations from the study mean. For each study, the effects of covariates age, sex, site, and total in-scanner motion were regressed out from the FC. Sensitivity analyses were conducted using higher resolution atlases^44^ describing 114 and 219 cortical regions (data shown in Supplemental Results), and without applying global mean correction (data shown in Supplemental Results).

### Polyconnectomic score

Figure 1 outlines the steps to compute polyconnectomic score (PCS), a method that leverages ‘connectome summary statistics’ (CSS). These statistics represent the strength and direction of the association between brain connections and the phenotype of interest (POI)^45^. CSS are typically represented by a symmetric matrix β of size *n* by *n*, where each element denotes the association strength of a connection between a pair of regions. CSS can be based on either a discovery dataset or a previously conducted independent study. In this work, we tested the PCS framework by deriving it from the CSS of a single study, or by aggregating CSS from multiple independent studies using a meta-analytic approach with a random-effects model^46^. The strength of an association between a connection and a POI is quantified using regression coefficients for scale variables or *t*-statistics and Cohen’s *d* for group contrasts. The PCS for an out-of-sample individual can then be computed as the weighted average of the CSS and the individual’s brain connectivity map:

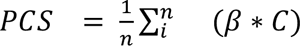

**Fig 1.**
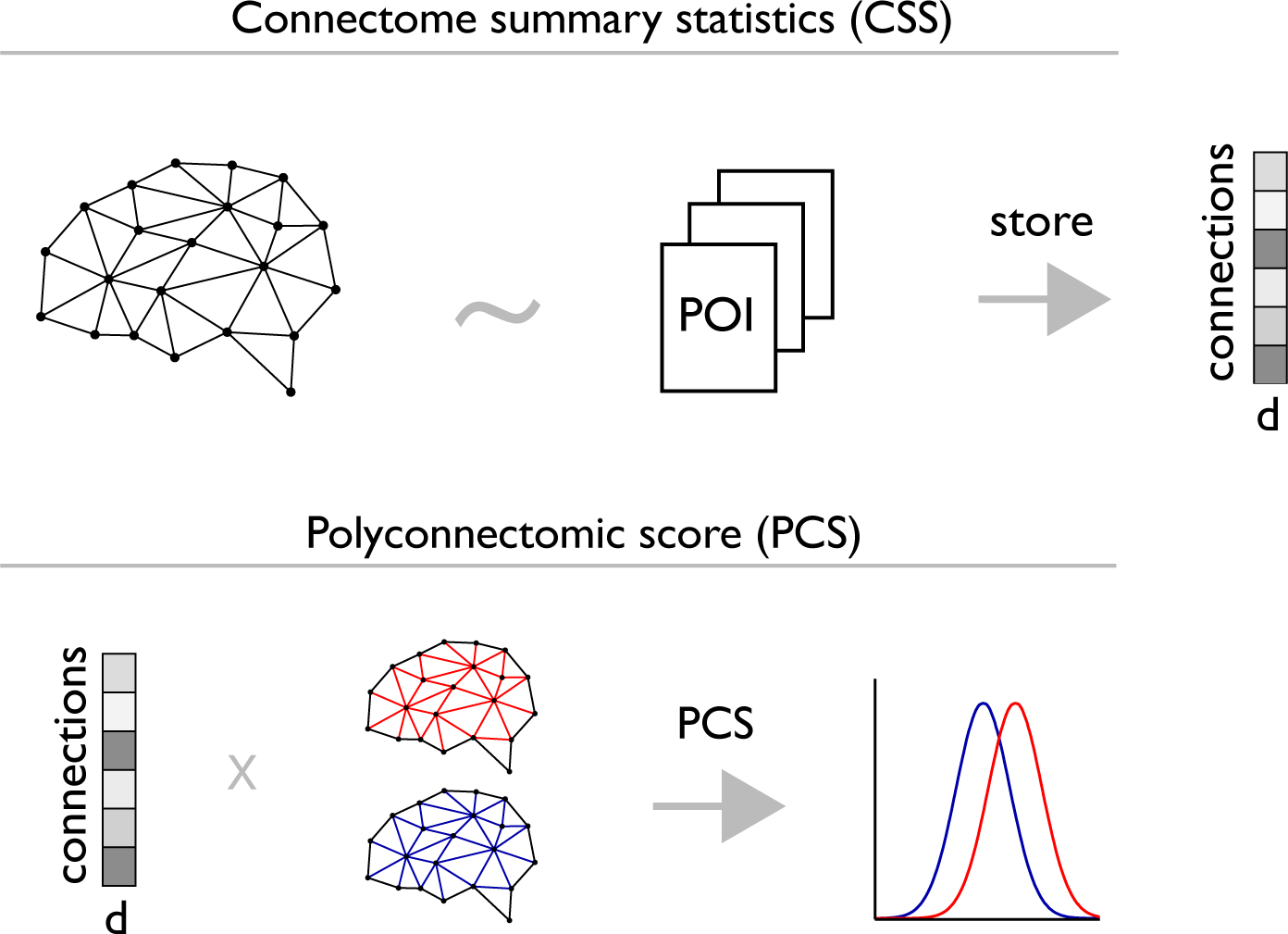
Computation of polyconnectomic score. The computation of the polyconnectomic score (PCS) relies on connectome summary statistics (CSS). These statistics represent the strength and direction of the association between brain connections and the phenotype of interest (POI), measured using regression coefficients for scaled variables or Cohen’s *d* for binary variables. CSS can be based on either a discovery dataset or a previously conducted independent study. The PCS for an out-of-sample individual can then be computed as the weighted average of the CSS and the individual’s brain connectivity map, capturing how closely a subject’s connectome resembles the brain signature associated with the POI. The efficacy of PCS is evaluated by comparing scores between cases and controls. *PCS*, polyconnectomic score; *POI*, phenotype of interest; *d*, Cohen’s d.

Here β is the CSS matrix and C is the connectivity map of the new subject. As such, PCS serves as a relative measure, attaining interpretative significance when compared with other subjects within the same sample.

### Statistical analyses

#### Simulations

We examined the theoretical predictive power of PCS by means of Monte Carlo simulations. A FC matrix was generated by sampling from a Gaussian distribution of size *n* by *n* (here, *n* = 68) with a mean of zero and standard deviation of 0.2, resulting in values ranging from −1 to 1. Studies were simulated with varying sample sizes (50, 100, 200, 500, 800, and 1600 subjects), and each study was partitioned into two equally-sized non-overlapping groups representing cases and controls. We generated a simulated contrast covering 10% of connections by sampling from a Gaussian distribution with a mean of zero and a standard deviation of 0.03, and subsequently applied this contrast to all cases. The simulated contrast resulted in a mixture of increased and decreased connectivity values in cases compared to controls, alterations equivalent to a distribution of Cohen’s *d* with mean zero and a standard deviation of 0.15. In each experiment, two studies of the same sample size were simulated. CSS were extracted from one study to compute PCS in the second study. We then evaluated the predictive power of PCS by estimating the Cohen’s *d* case-control differences in PCS. PCS was further evaluated using a logistic regression to classify each subject’s diagnosis and compute the area under the curve (*AUC*) of the receiver operating characteristic curve, with an *AUC* above 0.5 indicating a prediction better than random chance. The experiments were conducted in two scenarios: in the first scenario, cases from both studies received alterations on identical connections; in the second scenario, the alteration was applied in randomly different connections between studies. We conducted the experiment 1,000 times, considering both connectivity alteration scenarios, various sample sizes, and multiple *p*-value thresholds for the inclusion of connections from the CSS (ranging from 1 to 1 x 10^−5^).

#### Connectome summary statistics

We computed PCS using CSS representing FC differences between patients with neuropsychiatric disorders and controls. For the section on phenotypic prediction, we sourced CSS from the studies that had the largest patient samples available to us for ASD, SCZ, ADHD, and AD^26,29,31,33^. In the subsequent analyses, we conducted a meta-analysis using the CSS from four studies focusing on SCZ patients^25,30,32,33^ and four studies on ASD patients^26,28,29,33^ to compute PCS for SCZ (PCS-SCZ) and PCS for ASD (PCS-ASD), respectively. In each analysis, we ensured that the CSS were derived from studies independent of those where PCS was calculated. Table S1 provides the meta-analytic CSS for each disorder, allowing researchers to compute PCS for ASD, SCZ, ADHD, and AD in their own studies.

#### Phenotypic prediction

PCS was computed for patients with ASD, SCZ, ADHD, AD, and healthy controls. We assessed the ability of PCS to differentiate between patients and controls using Cohen’s *d*, obtaining *p*-values from a Student’s *t*-test and correcting for false discovery rate (FDR)^47^. The ability of PCS to classify individuals’ disease status more accurately than random chance was evaluated through a logistic regression with PCS as the predictor and diagnosis as the outcome variable. We computed the *AUC* to evaluate the model’s accuracy, using class weights to balance the differences in sample size between patient and control groups.

#### Patient stratification based on disease liability

We carried out analyses to investigate whether PCS can distinguish individuals based on their predisposition to psychosis. PCS-SCZ was computed in an independent study including patients with SCZ, SCA, BD, their first-degree relatives, and healthy controls^36^. PCS-SCZ levels were statistically compared across all groups using Cohen’s *d*, and *p*-values derived from Student’s *t*-test statistic (FDR-corrected).

#### Brain-behavior associations

We evaluated the utility of PCS in uncovering brain-behavior associations. We computed PCS-SCZ and PCS-ASD within the UKB cohort^37^. The relationship between PCS and cognitive measures, mental health factors, psychosocial questionnaires, and body measurements was analyzed. Pearson’s correlation coefficient was estimated for scaled variables, and Cohen’s *d* for categorical variables with *p*-values derived from Student’s *t*-test statistic (FDR-corrected). We replicated the analysis focusing on a subclinical population of subjects without clinical records of neuropsychiatric disorders or self-reported diagnoses (Supplemental Results).

## Results

### Evaluating PCS performance through simulation studies

In each iteration, we simulated two studies (Methods). The connectome summary statistics (CSS) from the first simulated study were extracted to compute polyconnectomic score (PCS) in the second study (PCS framework is illustrated in Figure 1). Simulations showed that when two studies presented different sets of altered connections, PCS revealed no significant differences between cases and controls (Figure S1). On the other hand, when identical connections were manipulated in cases from both studies, and the sample size was 100 individuals or more, PCS was higher in cases compared to controls (*p* < 0.05; Figure S1). In simulations with 100 individuals, group differences in PCS decreased when increasing the *p*-value thresholds for the inclusion of connections from the CSS, with Cohen’s *d* ranging from 0.52 (no *p*-value threshold) to 0.22 (*p*-value threshold < 5 x 10^−4^). For larger sample sizes, the predictive power of PCS declined when reaching Bonferroni correction. In simulations with 400 individuals, Cohen’s *d* for group differences in PCS started at 0.97 (no *p*-value threshold), peaked at 1.18 (*p*-value threshold < 0.01), and then declined, reaching 0.61 at Bonferroni correction (*p*-value threshold < 1 x 10^−5^). Only in simulations with 1,600 individuals, we found comparable group differences in PCS, regardless of whether all connections were included (*d* = 1.56) or Bonferroni correction was applied (*d* = 1.63).

### PCS capture brain patterns related to neuropsychiatric disorders

We evaluated the ability of PCS to identify ASD-associated brain connectivity patterns and differentiate patients from controls (Methods; Figure 1). ASD patients exhibited significantly higher PCS-ASD levels compared to controls (*d* = 0.45, *p* = 3 x 10^−11^, FDR-corrected), and using PCS-ASD led to a classification of individuals’ disease status more accurate than would expected by random chance (*AUC* = 0.63). We further evaluated the reliability of PCS in detecting ASD-related brain signatures by incorporating two additional independent studies. Consistent with our initial findings, ASD patients in both studies displayed higher PCS-ASD compared to controls (*d* = 0.30 and 0.43, *p* = 2 x 10^−3^ and 1.2 x 10^−3^, FDR-corrected; *AUC* = 0.58 and 0.61, respectively; Figure 2A; Table S2).

**Figure 2.**
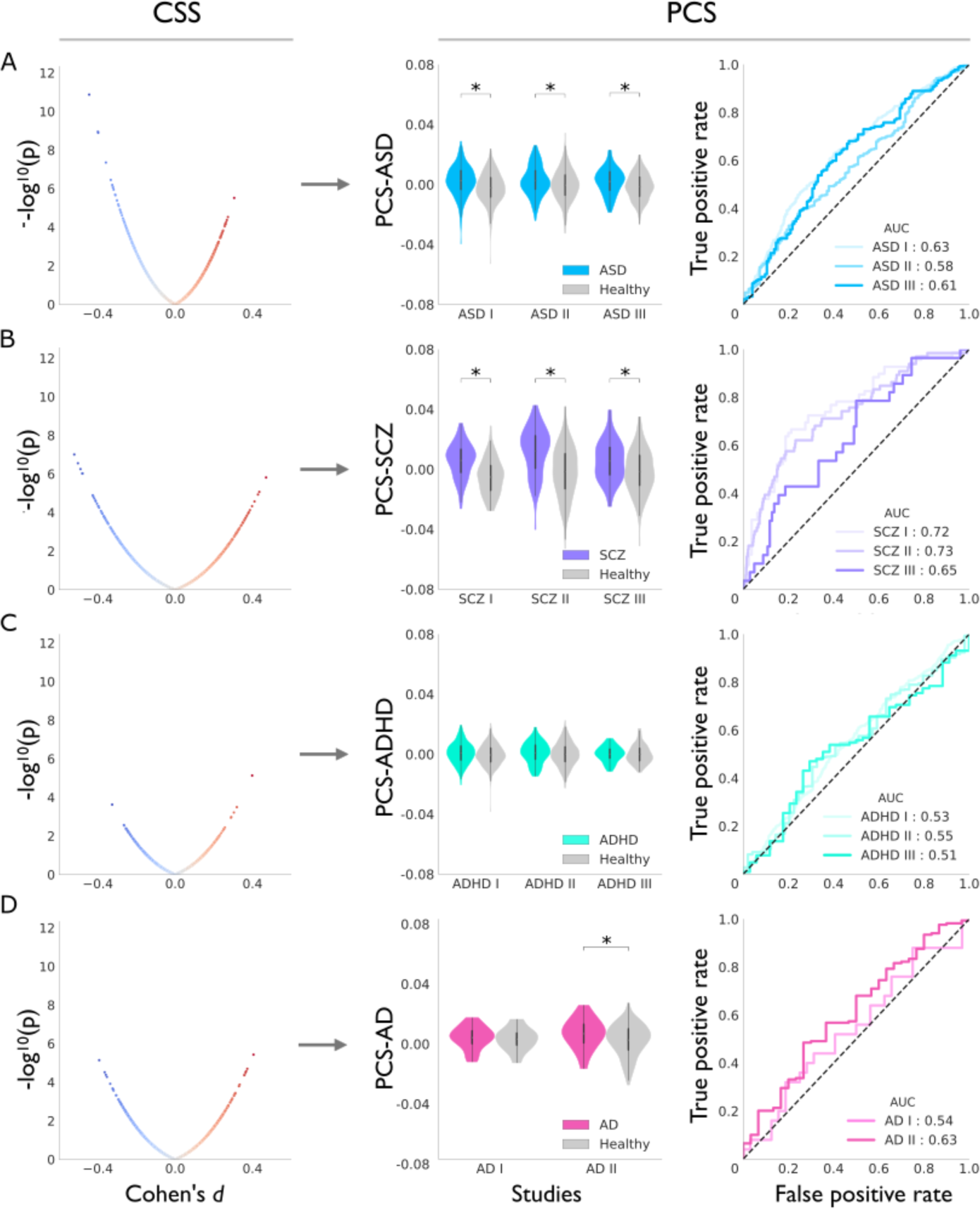
Polyconnectomic score for autism spectrum disorder, schizophrenia, attention deficit hyperactivity disorder, and Alzheimer’s disease. Connectome summary statistics (CSS) are estimated from a previously conducted study (left column). These statistics quantify the strength of the association (x-axis) and the level of significance (y-axis) for each brain connection in relation to (*A*) autism spectrum disorder, (*B*) schizophrenia, (*C*) attention deficit hyperactivity disorder, and (*D*) Alzheimer’s disease. Blue and red dots represent connections with decreased and increased functional connectivity in patients compared to controls, respectively. These CSS are used to calculate the polyconnectomic score (PCS) in an independent study (middle column). In each study, we statistically compare PCS levels between patients and controls (y-axis; white dots indicate group means) to assess the method’s efficacy in capturing brain connectivity signatures linked to neuropsychiatric disorders. Asterisks denote studies where significant differences in PCS between groups were observed, as estimated by *t*-test statistics (FDR-corrected). Logistic regression analysis (right column) is used to evaluate the predictive power of PCS in classifying individual diagnoses by estimating the area under the receiver operating characteristic curve (*AUC*; x-axis for false positive rate, y-axis for true positive rate). A dotted line at an *AUC* of 0.5 corresponds to random guessing. *CSS*, connectome summary statistics; *PCS*, polyconnectomic score; *ASD*, autism spectrum disorder; *SCZ*, schizophrenia; *ADHD*, attention deficit hyperactivity disorder; *AD*, Alzheimer’s disease; *AUC*, area under the curve of the receiver operating characteristic curve.

We expanded our analyses to compute PCS for SCZ, ADHD, and AD (Table S2). In three studies, individuals diagnosed with SCZ displayed elevated PCS-SCZ relative to controls ([min, max]: *d* = [0.54, 0.87], *p* < 0.05, FDR-corrected; *AUC* = [0.65, 0.73]; Figure 2B). Individuals with ADHD showed no differences in PCS-ADHD compared to controls (*p* > 0.05, FDR-corrected; Figure 2C). In one study, AD patients presented elevated PCS-AD compared to controls (*d* = 0.48, *p* = 0.034, FDR-corrected; *AUC* = 0.63), while no such difference was found in the other study (*p* > 0.05, FDR-corrected; Figure 2D). Differences in PCS between groups remained consistent under various conditions, including evaluations against a null model that permuted the effect sizes from the CSS (Table S2), computations of PCS using a meta-analytical CSS (Figure 3 and Table S3), utilizing higher-resolution atlases or without applying global mean correction (Supplemental Results).

**Figure 3.**
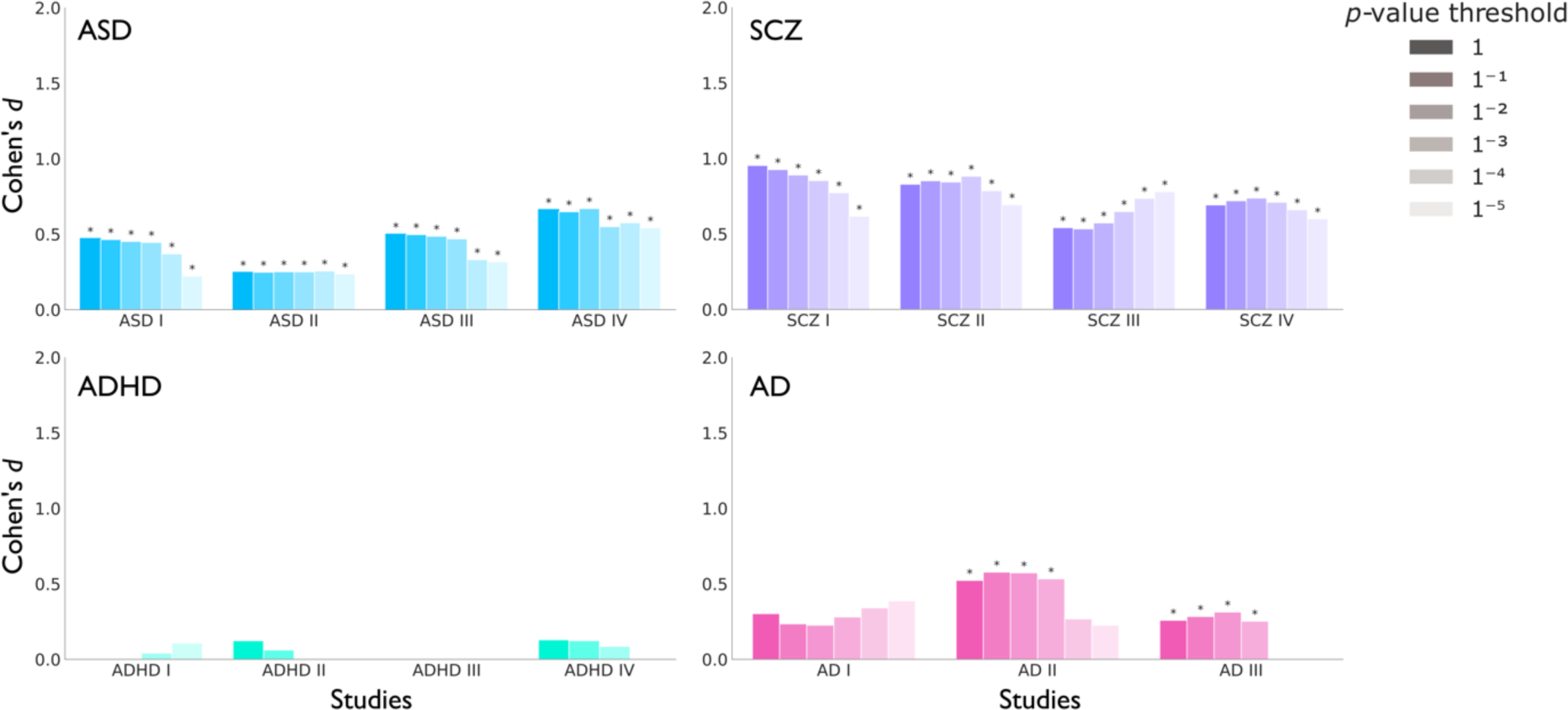
Computation of polyconnectomic score using meta-analytic summary statistics. A leave-one-out meta-analysis is conducted to derive robust connectome summary statistics (CSS) for calculating the polyconnectomic score (PCS) in independent studies. Within each study (x-axis), differences in PCS levels between patients and controls are estimated using Cohen’s *d* (y-axis). Connections from the CSS are thresholded based on *p*-value significance levels, ranging from no threshold to approximately Bonferroni correction (*p*-value threshold < 1 x 10^−5^). Asterisks indicate studies where significant differences in PCS between patients and controls are observed, as estimated by *t*-test statistics (FDR-corrected). *PCS*, polyconnectomic score; *ASD*, autism spectrum disorder; *SCZ*, schizophrenia; *ADHD*, attention deficit hyperactivity disorder; *AD*, Alzheimer’s disease.

### PCS stratifies individuals across the psychosis continuum

We investigated the ability of PCS to differentiate individuals based on their predisposition to psychosis by computing PCS-SCZ for patients with SCZ, first-degree relatives of SCZ patients, and healthy controls (Methods). Analysis revealed that SCZ patients exhibited higher PCS-SCZ compared to both their first-degree relatives (*d* = 0.42, *p* = 4 x 10^−3^, FDR-corrected; Figure S7) and healthy controls (*d* = 0.77, *p* = 1 x 10^−6^, FDR-corrected). First-degree relatives of SCZ patients also presented significantly elevated PCS-SCZ compared to healthy controls (*d* = 0.34, *p* = 0.034, FDR-corrected).

We expanded our analysis to the complete psychosis continuum, including patients with schizoaffective disorder (SCA) and bipolar disorder (BD), as well as their first-degree relatives (Figure 4). SCZ patients showed the largest difference in PCS-SCZ compared to healthy controls (*d* = 0.77), followed by patients with SCA (*d* = 0.66, p = 7.4 x 10^−4^, FDR-corrected) and BD (*d* = 0.57, *p* = 1.6 x 10^−3^, FDR-corrected). No statistical differences were observed in PCS-SCZ among SCZ, SCA, and BD patients (*p* > 0.05, FDR-corrected). Within individual disorders, patients with SCA exhibited PCS-SCZ levels that were nominally higher than those of their first-degree relatives (*d* = 0.31, *p* = 0.12, FDR-corrected), and first-degree relatives also showed a slight increase in PCS-SCZ compared to healthy controls (*d* = 0.35, *p* = 0.11, FDR-corrected), although these effects did not reach significance after correction for multiple comparisons. BD patients presented a significant increase in PCS-SCZ compared to their first-degree relatives (*d* = 0.59, *p* = 1.6 x 10^−3^, FDR-corrected), while first-degree relatives of BD patients showed no differences with healthy controls (*p* > 0.05, FDR-corrected).

**Figure 4.**
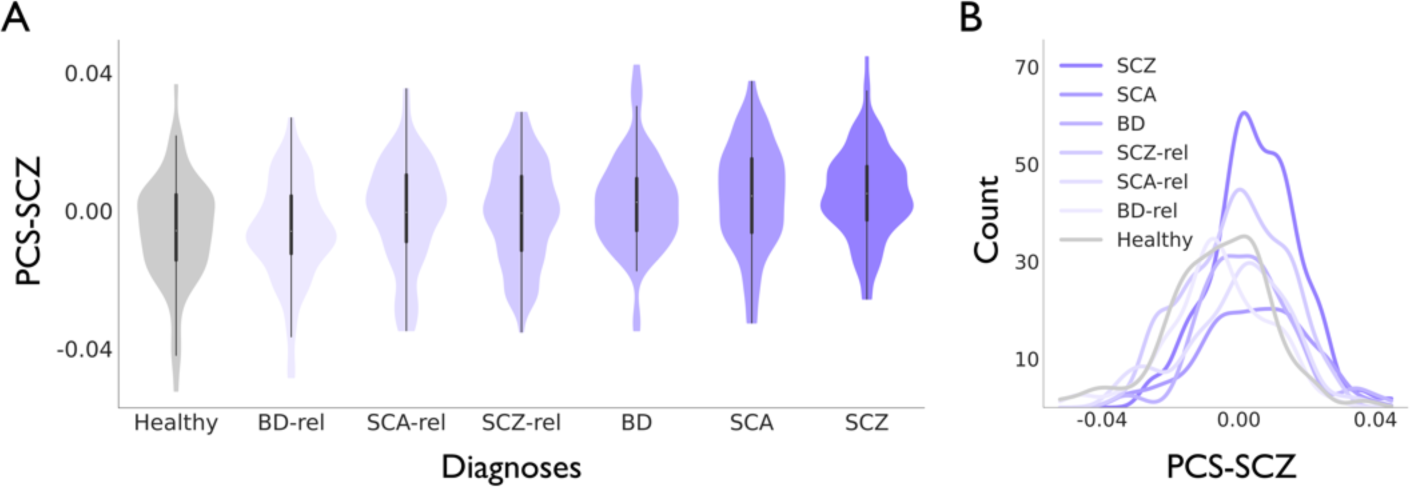
Using polyconnectomic score for schizophrenia to stratify individuals across the psychosis continuum. The polyconnectomic score (PCS) for schizophrenia (SCZ) is computed across the psychosis continuum, including patients with SCZ, schizoaffective disorder (SCA), and psychotic bipolar disorder (BD), as well as first-degree relatives from each group, and healthy controls. (*A*) Violin plots display the distribution of PCS-SCZ (y-axis) for each group (x-axis; white dot denotes the mean). (*B*) A histogram shows the frequency count (y-axis) of PCS-SCZ values (x-axis) among individuals in each group. Patients with SCZ present the largest differences in PCS-SCZ compared to healthy controls (*d* = 0.77), followed by SCA (*d* = 0.66) and BD (*d* = 0.57). *SCZ*, schizophrenia; *SCA*, schizoaffective disorder; *BD*, bipolar disorder; *SCZ-rel*, first-degree relatives of schizophrenia patients; *SCA-rel*, first-degree relatives of schizoaffective disorder patients; *BD-rel*, first-degree relatives of bipolar disorder patients; *PCS-SCZ*, polyconnectomic score for schizophrenia.

### PCS detects brain-behavior associations

We further assessed the relationship between PCS and clinical as well as behavioral measures in the UK Biobank (UKB; Methods). Individuals exhibiting higher PCS-SCZ displayed lower fluid intelligence (*r* = −0.037, *p* = 1.1 x 10^−5^, FDR-corrected) and slower reaction times (*r* = 0.033, *p* = 5.7 x 10^−5^, FDR-corrected; Figure 5). No specific effects on cognition were found for PCS-ASD (Figure S8). Regarding mental health indicators, elevated PCS-SCZ was correlated with a higher likelihood of experiencing nervous feelings (*d* = 0.12, *p* = 2.8 x 10^−9^, FDR-corrected), increased neuroticism scores (*r* = 0.031, *p* = 1.5 x 10^−5^, FDR-corrected), and decreased propensity for risk-taking (*d* = −0.09, *p* = 1.8 x 10^−7^, FDR-corrected), among other aspects (complete results in Table S4). Subjects with higher PCS-ASD were likely to have more consultations with a psychiatrist for issues related to nerves, anxiety, tension, or depression (*d* = 0.08, *p* = 4.7 x 10^−3^, FDR-corrected). Our analysis also uncovered potential links between PCS and measures of well-being. We observed a negative correlation between happiness and both PCS-SCZ (*r* = −0.023, *p* = 6.4 x 10^−4^, FDR-corrected) and PCS-ASD (*r* = −0.026, *p* = 9.1 x 10^−4^, FDR-corrected). Individuals with higher PCS-SCZ reported lower levels of job satisfaction (*r* = −0.047, *p* = 1.6 x 10^−12^, FDR-corrected) and health satisfaction (*r* = −0.032, *p* = 1.1 x 10^−6^, FDR-corrected), whereas individuals with elevated PCS-ASD reported reduced levels of friendship satisfaction (*r* = −0.017, *p* = 0.033, FDR-corrected) and family satisfaction (*r* = −0.021, *p* = 7.1 x 10^−3^, FDR-corrected). These findings were consistent when focusing on healthy subjects without any neuropsychiatric diagnosis (Table S5).

**Figure 5.**
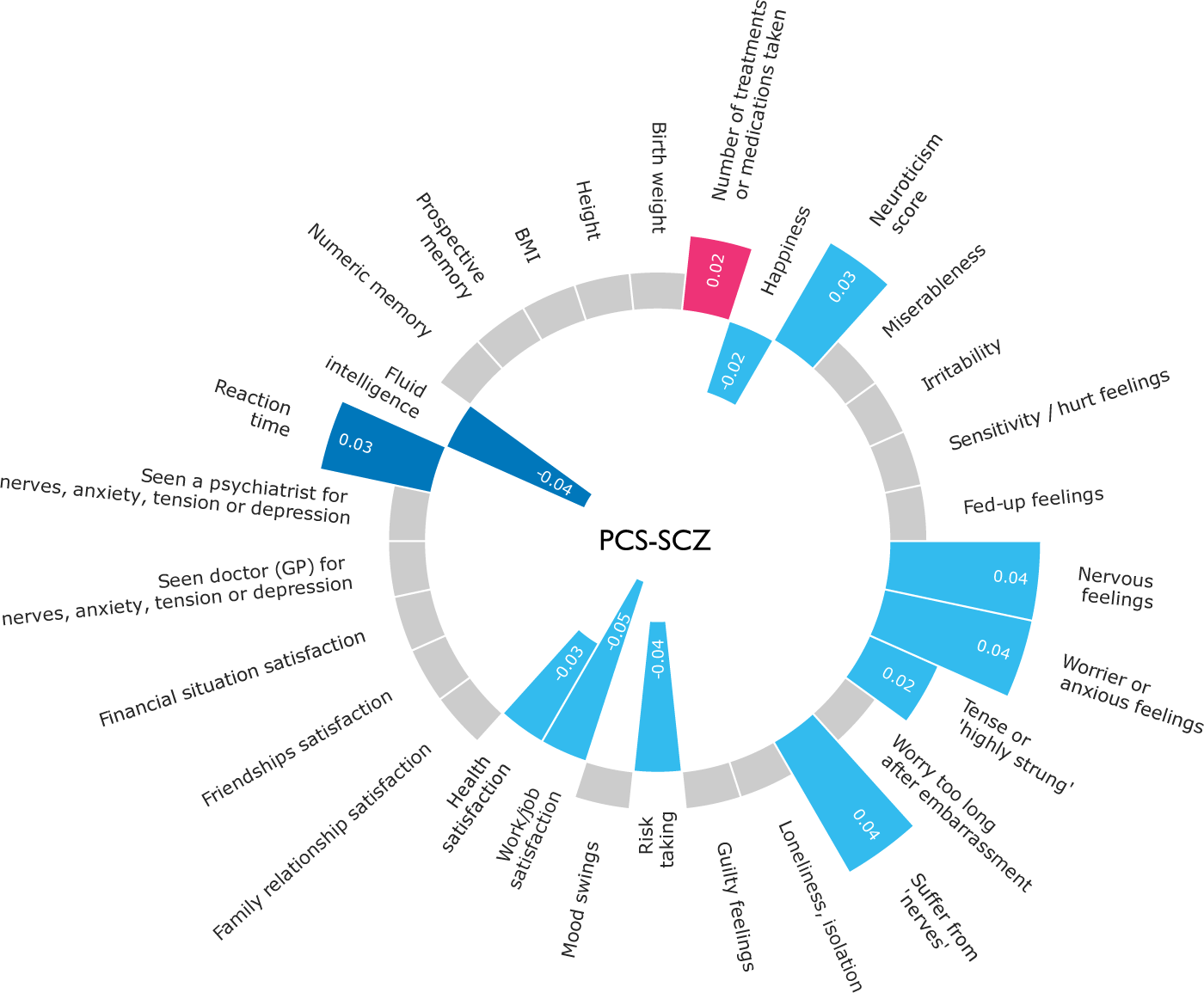
Brain-behavior correlations using polyconnectomic score for schizophrenia. A circle plot illustrates the association between the polyconnectomic score for schizophrenia (PCS-SCZ) and measures of cognition (dark blue), mental health (light blue), and self-reported medical conditions (pink), based on data from the UK Biobank. Pearson’s correlation coefficients are displayed only for associations that remain significant after FDR correction (gray indicates non-significant effects). Dichotomous variables measured with Cohen’s *d* are converted to Pearson’s correlation coefficient for visualization purposes. Elevated levels of PCS-SCZ are associated with reduced cognitive performance, increased neuroticism, and a higher incidence of mental health complaints. Similar associations are observed in subclinical populations. *PCS-SCZ*, polyconnectomic score for schizophrenia.

## Discussion

We investigated the ability of PCS to capture and quantify disorder-related brain signatures in individual connectomes. Our findings suggest that PCS has several valuable applications, including distinguishing populations with neuropsychiatric disorders, stratifying patients by disease risk, and uncovering links between brain connectivity and behavior.

PCS associated with neuropsychiatric disorders were generally higher in patients compared to controls. SCZ patients in particular showed elevated PCS-SCZ than controls across three different datasets ([min, max]: *d* = [0.54, 0.87]; Figure 2B and Table S2), supporting the hypothesis of dysconnectivity as a basis for the neuropathology of SCZ^48,49^. The mean effect of PCS-SCZ across studies (*d* = 0.74) exceeded the largest effect sizes of previously reported brain alterations in SCZ, such as cortical thickness thinning (*d* = −0.54)^50^, reductions in total gray matter (*d* = - 0.58)^51^ and thalamus volume (*d* = −0.68)^51^, enlargement of the third ventricle (*d* = 0.60)^51^, and alteration in overall white matter microstructure (*d* = −0.42)^52^. ASD patients similarly showed higher PCS-ASD than controls in three studies (*d* = [0.30, 0.45]; Figure 2A and Table S2) with a mean effect size of 0.39, an effect comparable to the largest effect sizes reported for whole brain thickness in ASD patients (*d* = 0.41)^53^. Furthermore, larger group differences in PCS were generally found as a result of employing a meta-analytic CSS for the computation of PCS (Figure 3 and Table S3). These results support the utility of PCS as a valuable brain metric for distinguishing patient groups with neuropsychiatric disorders from control populations.

PCS differentiated patients from controls more robustly when all connections were included in the analysis, compared to when only connections that remained significant after Bonferroni correction were incorporated. Both our simulations (Figure S1) and empirical analyses (Figure 3) showed that connections containing relevant information about the phenotype under investigation often were discarded in traditional neuroimaging studies after controlling for multiple comparisons. Such findings are consistent with existing evidence suggesting that effects found in spatially localized sets of connections may constitute only a small fraction of a broader global brain involvement^14–16^. In line with this evidence, differences in PCS between patients and controls exceeded those in global FC, potentially due to the inclusion of the entire connectome combined with the disease-specific circuitry information (Figure S4 and Figure S5).

A crucial question regarding PCS is whether this method captures connectivity patterns specific to the disease of interest or whether it represents a general alteration common across disorders. Our data indicate that PCS-ASD levels are higher in ASD patients compared to controls, an effect generally not observed when comparing PCS-ASD levels between individuals with other neuropsychiatric disorders and controls (Figure S3). These observations suggest that PCS was able to capture disease-specific brain predispositions, rather than a general cross-disorder vulnerability^19–22^, which is critical for developing connectomic markers with potential clinical applications^3^. However, PCS was not always disease-specific.

Patients with BD, ASD, and major depressive disorder also exhibited elevated PCS-SCZ compared to controls (Figure S3). Additional analyses indicated that PCS-SCZ appears to identify brain signatures related to psychosis liability, not exclusively specific to SCZ. Across the psychosis continuum, SCZ patients presented the highest PCS-SCZ levels, followed by SCA and BD patients, as well as their respective first-degree relatives (Figure 4), suggesting an overlap in the functional brain circuitry alterations across psychotic disorders^22,54^.

PCS shows potential in uncovering associations between brain connectivity and cognitive measures, mental health factors, psychosocial questionnaires, and body measurements. Individuals with connectomes resembling those typically seen in SCZ exhibited reduced cognitive performance, echoing the well-established relationship between SCZ and intelligence (Figure 5)^55–58^. Additionally, we observed significant correlations of PCS-SCZ and PCS-ASD with neuroticism. This observation aligns with existing research identifying links between neuroticism and both SCZ and ASD at the behavioral^59,60^ and genetic levels^61–63^. Given that neuroticism and intelligence are associated with a general factor of psychopathology (*p*-factor)^64^, it is plausible that PCS-SCZ could serve as a broader marker for transdiagnostic psychopathology. In terms of physical measurements, elevated PCS-ASD was associated with a lower body mass index (Figure S8). A negative association between male children with ASD and body mass index has been previously reported^65^, although other studies report a positive association^66,67^. These findings demonstrate the utility of PCS for detecting links between brain connectivity and behavior, contributing to a deeper understanding of the neural underpinnings of clinical traits.

PCS sets itself apart from existing methods such as examining graph metrics^1^, network-based statistics^68^, regional vulnerability index^69^, and connectome-based predictive modeling^70^. One key strength of PCS is the inclusion of whole-brain connectivity information to provide an interpretable metric that quantifies the resemblance of a given connectome to the neural circuitry associated with a phenotype. PCS is not constrained by the topological organization of brain circuits related to a phenotype, offering a comprehensive and unbiased view on the implicated brain connections. PCS is also designed to take into account both the strength of individual connections and their similarity to the phenotype-associated neural circuitry, maximizing the information utilized to detect relevant brain signatures. Additionally, PCS is a computationally inexpensive method and does not require data normalization across subjects, reducing the risk of data leakage between patient and control groups. Previous studies have used frameworks similar to PCS for analyzing brain functional activation^71^ and connectivity^72^, providing compelling evidence of the predictive power to detect individual differences in cognitive performance. Building upon this work, our study replicates these findings in the UKB (Figure S6), and further extends the application of PCS to both healthy and diseased connectomes across multiple independent studies.

There are several limitations that should be noted. Similar to polygenic score^23^, PCS serves as a relative measure and its interpretation becomes meaningful only within the context of the same sample. The development of an absolute scale is crucial to enhance its clinical utility. PCS do not consider potential interactions within the brain, treating each connection independently. A model that accounts for the autocorrelated structure of the brain could improve the predictive power of neuroimaging markers^71,73,74^, although this is not always the case^72^. Previous studies have highlighted the potential of connectivity markers in predicting the progression of brain disorders^75,76^. While PCS distinguished groups of individuals based on psychosis risk in cross-sectional analyses, the ability of PCS to predict disease onset in longitudinal studies remains to be determined. Furthermore, PCS showed limited effectiveness in distinguishing ADHD patients from controls. This result could be due to a pronounced disease heterogeneity across ADHD studies (Figure S2), potentially reducing the predictive power of PCS in identifying disorder-related brain patterns in diverse patient groups.

PCS provides a way to address the challenges inherent in network neuroimaging studies. Using PCS to integrate information from the entire brain into a single score has the potential to create robust endophenotypes while boosting statistical power. We show evidence that PCS can aid researchers to identify subjects with neuropsychiatric disorders, stratify individuals based on disease risk, and uncover brain-behavior associations. PCS stands as a promising tool for deepening our understanding of both healthy and diseased brains, with a wide range of potential applications in the field of neuroscience.

## Data availability

COBRE is available from http://schizconnect.org. CNP is available from OpenNeuro (http://openneuro.org, accession number ds000030). BSNIP is obtained from the NIMH Data Archive (https://nda.nih.gov; NDAR ID: 2274, respectively). SRPBS is available at https://bicr-resource.atr.jp/srpbs1600. HBN is available at http://fcon_1000.projects.nitrc.org/indi/cmi_healthy_brain_network. NKI-Enhanced is obtained from http://fcon_1000.projects.nitrc.org/indi/enhanced/. ADNI2/GO, and ADNI 3 are available from https://ida.loni.usc.edu. OASIS3 is available at www.oasis-brains.org. The UKB MRI data is available at https://www.ukbiobank.ac.uk.

## Author contributions

M.P.v.d.H. and I.L. conceived the project. I.L. analyzed the data. M.P.v.d.H, and I.L. wrote the manuscript. K.H., L.G.S., M.G., J.R., T.K., and U.D., provided expertise and feedback on the text and analyses.

## Acknowledgments

The research of M.P.v.d.H. was funded by a ERC Consolidator Grant (101001062) from the European Research Council (ERC).

This research has been conducted using the UK Biobank resource under application 16406. We thank the numerous participants, researchers, and staff from many studies who collected and contributed to the data, especially to Jeanne Savage, Siemon de Lange, and Elleke Tissink for their contributions to the development of the pipelines for the UK Biobank data utilized in this study.

Analyses were carried out on the Genetic Cluster Computer hosted by the Dutch National computing and Networking Services SURFsara and financed by the Netherlands Organization for Scientific Research (NWO: 480-05-003), the VU University (Amsterdam, The Netherlands) and the Dutch Brain Foundation.

COBRE data were downloaded from the Collaborative Informatics and Neuroimaging Suite Data Exchange tool (COINS; http://coins.mrn.org/dx), and data collection was performed at the Mind Research Network and funded by the Center of Biomedical Research Excellence (COBRE) grant 5P20RR021938/P20GM103472 from the NIH to V. Calhoun.

Data and/or research tools used in the preparation of this manuscript were obtained from the National Institute of Mental Health (NIMH) Data Archive (NDA). NDA is a collaborative informatics system created by the National Institutes of Health to provide a national resource to support and accelerate research in mental health. Dataset identifier(s) is 2274. This manuscript reflects the views of the authors and may not reflect the opinions or views of the NIH or of the Submitters submitting original data to NDA.

Data collection and sharing for SRPBS was provided by the DecNef Department at the Advanced Telecommunication Research Institute International, Kyoto, Japan.

Data collection and sharing for this project was funded by the Alzheimer’s Disease Neuroimaging Initiative (ADNI) (National Institutes of Health Grant U01 AG024904) and DOD ADNI (Department of Defense award number W81XWH-12-2-0012). ADNI is funded by the National Institute on Aging, the National Institute of Biomedical Imaging and Bioengineering, and through generous contributions from the following: AbbVie, Alzheimer’s Association; Alzheimer’s Drug Discovery Foundation; Araclon Biotech; BioClinica, Inc.; Biogen; Bristol-Myers Squibb Company; CereSpir, Inc.; Cogstate; Eisai Inc.; Elan Pharmaceuticals, Inc.; Eli Lilly and Company; EuroImmun;

F. Hoffmann-La Roche Ltd and its affiliated company Genentech, Inc.; Fujirebio; GE Healthcare; IXICO Ltd.; Janssen Alzheimer Immunotherapy Research & Development, LLC.; Johnson & Johnson Pharmaceutical Research & Development LLC.; Lumosity; Lundbeck; Merck & Co., Inc.; Meso Scale Diagnostics, LLC.; NeuroRx Research; Neurotrack Technologies; Novartis Pharmaceuticals Corporation; Pfizer Inc.; Piramal Imaging; Servier; Takeda Pharmaceutical Company; and Transition Therapeutics. The Canadian Institutes of Health Research is providing funds to support ADNI clinical sites in Canada. Private sector contributions are facilitated by the Foundation for the National Institutes of Health (www.fnih.org). The grantee organization is the Northern California Institute for Research and Education, and the study is coordinated by the Alzheimer’s Therapeutic Research Institute at the University of Southern California. ADNI data are disseminated by the Laboratory for NeuroImaging at the University of Southern California.

Data were provided [in part] by OASIS (T. Benzinger, D. Marcus, J. Morris; NIH P50 AG00561, P30 NS09857781, P01 AG026276, P01 AG003991, R01 AG043434, UL1 TR000448, R01 EB009352. AV-45 doses were provided by Avid Radiopharmaceuticals, a wholly owned subsidiary of Eli Lilly).

We acknowledge the contribution of data made available through SchizConnect (http://schizconnect.org; funded by NIMH [grant 1U01 MH097435]), NIMH Data Archive (https://nda.nih.gov), OpenNeuro (http://openneuro.org), and XNAT Central (http://central.xnat.org).

The FOR 2107 cohort was funded by the German Research Foundation (DFG grants FOR2107 KI588/14-1, and KI588/14-2, and KI588/20-1, KI588/22-1 to Tilo Kircher, Marburg, Germany). Biosamples and corresponding data were sampled, processed and stored in the Marburg Biobank CBBMR.

